# SARS-CoV-2 nsp1 mediates broad inhibition of translation in mammals

**DOI:** 10.1101/2025.01.14.633005

**Authors:** Risako Gen, Amin Addetia, Daniel Asarnow, Young-Jun Park, Joel Quispe, Matthew C. Chan, Jack T. Brown, Jimin Lee, Melody G Campbell, Christopher P. Lapointe, David Veesler

**Affiliations:** Department of Biochemistry, University of Washington; Seattle, WA 98195, USA; Division of Basic Sciences, Fred Hutchinson Cancer Center, Seattle, WA, USA; Howard Hughes Medical Institute, University of Washington; Seattle, WA 98195, USA

## Abstract

SARS-CoV-2 nonstructural protein 1 (nsp1) promotes innate immune evasion by inhibiting host translation in human cells. However, the role of nsp1 in other host species remains elusive, especially in bats which are natural reservoirs of sarbecoviruses and possess a markedly different innate immune system than humans. Here, we reveal that SARS-CoV-2 nsp1 potently inhibits translation in bat cells from *Rhinolophus lepidus*, belonging to the same genus as known sarbecovirus reservoirs hosts. We determined a cryo-electron microscopy structure of SARS-CoV-2 nsp1 bound to the *Rhinolophus lepidus* 40S ribosome and show that it blocks the mRNA entry channel via targeting a highly conserved site among mammals. Accordingly, we found that nsp1 blocked protein translation in mammalian cell lines from several species, underscoring its broadly inhibitory activity and conserved role in numerous SARS-CoV-2 hosts. Our findings illuminate the arms race between coronaviruses and mammalian host immunity (including bats), providing a foundation for understanding the determinants of viral maintenance in bat hosts and spillovers.

## Introduction

Outbreaks of disease have plagued humans since the beginning of recorded history. Around 75% of human infectious diseases are thought to originate from spillover of non-human animal pathogens, also known as zoonosis^1^. Climate change, urbanization, and globalization all contribute to the increasing chances of new pathogens being introduced into the human population as a result of more frequent interactions between animals and humans^2^. Although many wild animals are natural reservoirs of pathogens, bats (Chiropteran order) are unusual in their ability to harbor more diverse viruses per animal species than other animals and they have been shown to act as reservoirs for multiple deadly viruses, such as coronaviruses, paramyxoviruses, and filoviruses^3^. Recent studies have suggested that unique characteristics of the bat innate immune system allow them to tolerate viruses that are highly pathogenic to humans and other animals^4–10^. However, little is known about the chiropteran immune responses to viral infection, partly due to the limited availability of tools and reagents to study bats^11–13^. Given the increasing number of spillover events of bat viruses to humans, there is a need to study the mechanisms of viral tolerance in bats to better predict and control future viral spillovers.

Within the past two decades, three coronaviruses with ancestral origins in bats have caused major outbreaks: severe acute respiratory syndrome coronavirus (SARS-CoV) in 2002, Middle East respiratory syndrome coronavirus (MERS-CoV) in 2012, and SARS-CoV-2 in 2019^14–18^,. SARS-CoV-2 is a positive-sense single stranded RNA virus that is part of the Betacoronavirus genus and Coronaviridae family^19^. The approximately 30 kb genome consists of 14 open reading frames (ORFs), of which ORF1a and ORF1b at the 5’ end are directly translated into two polyproteins by cellular ribosomes and processed to produce sixteen nonstructural proteins designated nsp1-nsp16^20^. These nsps work in unison and interfere with host cellular function in different ways to promote viral propagation.

Nsp1 is a major virulence factor of SARS-CoV-2 that antagonizes the host innate immune system through inhibition of host protein translation, blockage of mRNA nuclear export, promotion of host mRNA cleavage, and depletion of key host signaling factors^21–23^. Translation inhibition occurs through binding of the nsp1 C-terminal domain (CTD) to the mRNA entry channel within the human small ribosomal subunit (40S), sterically blocking host mRNA from entering the ribosome^21,24,25^. Whereas it leads to suppression of host mRNA translation, including the interferon response, viral mRNAs evade this nsp1-mediated translation inhibition through an unknown mechanism thought to include key interactions between the nsp1 N-terminal domain (NTD) and the SARS-CoV-2 mRNA 5’ untranslated region (UTR)^26,27^. Nsp1 mediates similar biological functions across alpha- and betacoronaviruses^28–30^, and three beta-CoV nsp1s were shown to inhibit translation through its binding to the human 40S ribosome^31,32^, highlighting its importance as a conserved virulence factor.

Beyond its direct impact on the global human population, SARS-CoV-2 infections have been reported in many other animals such as bats, dogs, cats, minks, and deers, showcasing its broad host tropism^33–36^. While viral entry into host cells relies on receptor engagement, the success of viral replication is dependent on the virus’s ability to hijack the infected host cell, as illustrated by the fact that although pig ACE2 supports spike-mediated cell entry, pigs are refractory to SARS-CoV-2 infection^37–40^. These results underscore the importance of identifying key viral determinants of successful SARS-CoV-2 replication in host animals. Given the poor understanding of the bat innate immune system and the marked differences from the human counterpart that emerged from prior studies, the virus-host dynamics leading to reservoir maintenance and spillovers remain elusive.

We set out to understand the impact of SARS-CoV-2 nsp1 on the innate immune response of bats and other mammals to get deeper insight into nsp1 function across diverse species. We used a *Rhinolophus lepidus* cell line to probe nsp1 function within bats as Rhinolophus horseshoe bats act as natural reservoirs of SARS-CoV-2 related coronaviruses^41,42^. Here, we unveil the architecture of the *Rhinolophus* 40S bat ribosome, demonstrate the broad translation inhibition activity mediated by nsp1 in bats and other mammalian species, and reveal a conserved binding mechanism of nsp1 to the bat ribosome. Our findings provide a foundation for understanding the determinants of nsp1-induced host innate immune antagonism and spillovers.

### SARS-CoV-2 nsp1 inhibits translation in *Rhinolophus* bat cells

To evaluate if SARS-CoV-2 nsp1 is functional in bat cells, we investigated nsp1-mediated inhibition of translation in a *Rhinolophus lepidus* kidney (Rhileki) cell line, which we selected as a model of sarbecovirus target cells. To quantify the impact of nsp1 activity on protein synthesis, we developed a fluorescence-based translation assay involving a GFP reporter mRNA and an nsp1 mRNA (**Fig. 1A**). The mRNA constructs are capped at the 5’ end and harbor a 80-nt poly-A tail at the 3’ end to mimic mature mRNAs, and contain N1-methyl pseudouridines (instead of uridines) as it was shown to dampen innate immune activation in humans cells^43^. We chose to include a synthetic non-viral 5’UTR for the GFP-encoding mRNA whereas the nsp1-encoding mRNA possesses the SARS-CoV-2 5’ UTR sequence to overcome nsp1-mediated translation inhibition^26,31^. Transfection of the GFP mRNA in both Rhileki and HEK293T cells led to robust expression observed by live-cell fluorescence imaging (**Fig 1B**). Co-transfection of the GFP and wildtype nsp1 mRNAs reduced fluorescence in an nsp1 mRNA dose-dependent manner with 75% reduction of GFP expression achieved with 0.01 ng of nsp1 mRNA and no detectable fluorescence remained with transfection of 1 ng of nsp1 mRNA in both cell lines (**Fig 1C**). Furthermore, the previously described nsp1 K164A/H165A double mutant^44^, which is unable to inhibit human ribosome-mediated translation, had no effect on GFP translation in Rhileki cells (**Fig 1C and S1**). These results indicate that nsp1 inhibits translation in both human and bat cells, most likely through a conserved mechanism involving interactions mediated by the nsp1 C-terminal domain, including residues K164 and H165.

**Fig 1.**
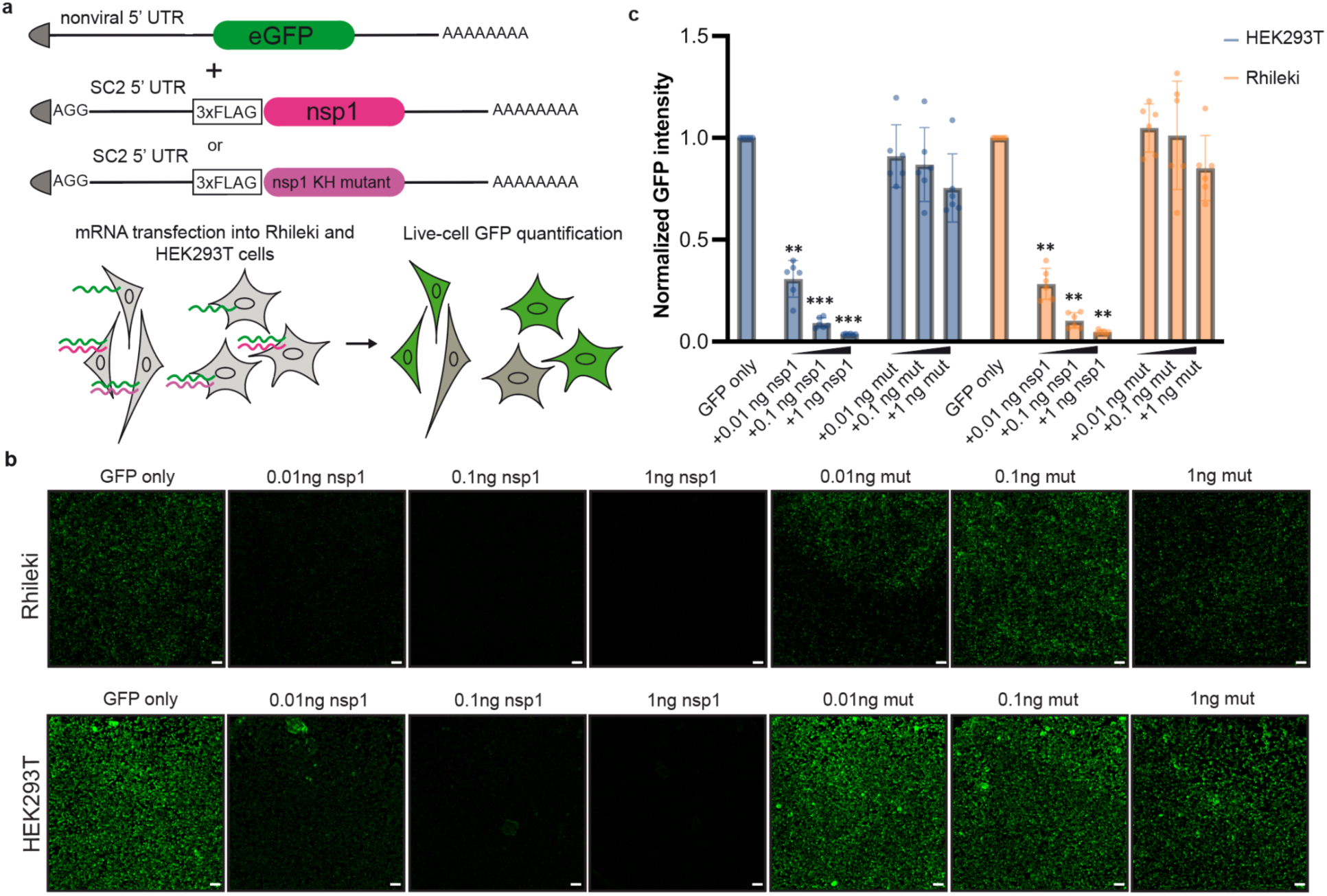
SARS-CoV-2 nsp1 suppresses translation in human (HEK293T) and Rhileki (bat) cells. (A) mRNA construct design and schematic of the fluorescence-based translation assay. 200 ng of eGFP-encoding mRNA was transfected with 0.01-1 ng of a wildtype or mutant nsp1 mRNA to enable visualization and quantification of translation inhibition. mRNAs are all 5’ capped (shown by the triangular grey shapes) and have poly(A) tails at the 3’ end. SC2: SARS-CoV-2. 5’-UTR: 5’ untranslated region.

3xFLAG: triple flag tag. nsp1 KH mutant: nsp1 K164A/H165A double mutant. (B) Live-cell fluorescence imaging of GFP expression in the presence (or absence) of varying amounts of wildtype nsp1 mRNA or of the K164A/H165A double mutant (mut) mRNA using Rhileki and HEK293T cells. Scale bar: 100 µm. See Fig S1 for GFP expression levels. (C) Quantification of the nsp1-mediated dose-dependent inhibition of translation based on normalized GFP fluorescence intensity across the entire field of view. Bars represent the mean of six biological replicates shown as individual data points with error bars showing standard deviation. Each biological replicate is a mean of six technical replicates. Transfections with wildtype and mutant nsp1 mRNAs were compared to the GFP-only control using one-way ANOVA and follow-up Dunnet’s T3 multiple comparisons tests. P-values reported are adjusted for multiplicity. **: p<0.01, ***: p<0.005.

### Architecture of the *Rhinolophus lepidus* bat ribosome

Although prior studies have provided a wealth of information on several prokaryotic and eukaryotic ribosomes^45–52^, the architecture and composition of the Chiropteran ribosome remain elusive. To address this knowledge gap and understand the mechanism of action of nsp1 in bats, we sought to purify ribosomes from the Rhileki cell line. Fractionation of Rhileki cell lysates through a 30% sucrose cushion (to pellet the ribosomes) followed by purification using a 11-25% sucrose density gradient enabled the isolation of pure 40S ribosomal subunits (**Fig S2**). The purified 40S subunits were subsequently incubated with recombinantly produced SARS-CoV-2 nsp1 and human eIF1 at a 1:5:5 molar ratio before sample vitrification for structural analysis. We used single particle cryo-EM to determine a structure of the SARS-CoV-2 nsp1-bound 40S Rhileki ribosome at an overall resolution of 2.1 Å (**Table S1**). Subsequent local refinements improved map resolvability for the 40S body and head, yielding maps at 2.1 Å resolution with significant improvements in the dynamic regions of the 40S subunit such as the 40S head region (**Fig S3**). Recombinant human eIF1 was added to promote stabilization of the nsp1 N-terminal domain (NTD), as previously described^31^. Although the eIF1 density was partially resolved in the cryoEM map, the nsp1 NTD density was too weak to be modeled. Given that there is no genomic information for Rhileki, we sequenced all but one 40S protein subunits via transcriptomics and used Sanger sequencing of the 18S rRNA and ribosomal protein S10 to enable model building of the entire 40S subunit (**Data S1, Table S2**). Comparing the Rhileki 40S and human 40S ribosome structures reveals a remarkable architectural and compositional similarity (**Fig 2A**), concurring with the conservation of their ribosomal subunit sequences (97-100% amino acid sequence identity) and of their 18S rRNA (99.52% nucleotide sequence identity). We note that the 60S ribosomal subunit sequences are less conserved than that of the 40S when comparing Rhileki to human ribosomes, with the lowest sequence identity for RPL29 being 85.1% (**Data S2**). Our data show that the Rhileki 40S ribosome is strikingly similar to the human ribosome despite extensive evolutionary divergence between the Chiropteran and Mammalian orders, likely reflecting the strong selective pressure in maintaining the functionality of the machinery mediating protein translation.

**Figure 2.**
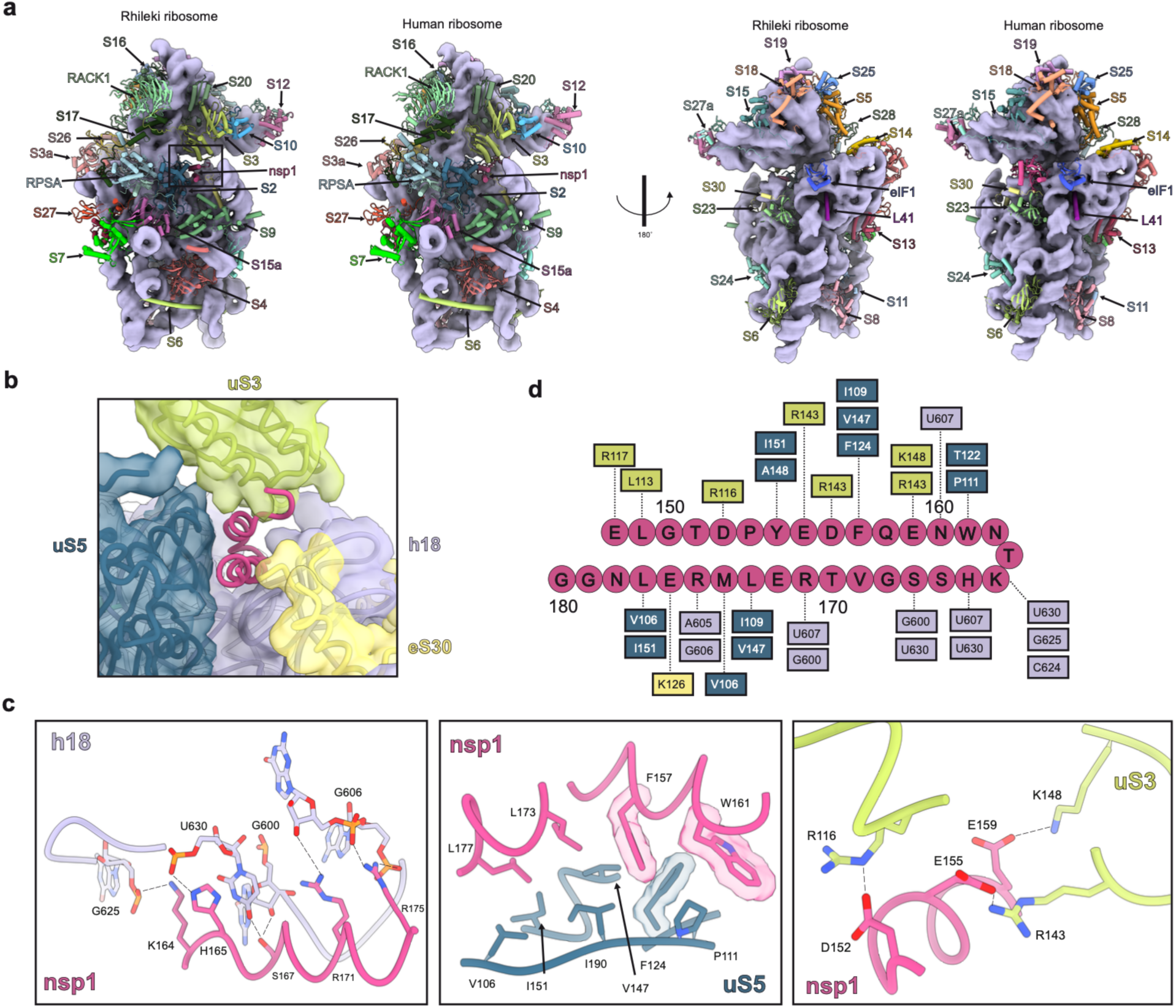
Architecture of the Rhileki bat 40S ribosome bound to SARS-CoV-2 nsp1. (A) Comparison of the 40S Rhileki ribosome and the human ribosome organization (PDB: 8PPK). Each ribosomal protein is colored distinctly and labeled whereas the 18S rRNA is rendered as a purple surface. The box showing the nsp1 binding interface is zoomed-in in panel B. (B) Closeup view of SARS-CoV-2 nsp1 (pink) within the mRNA entry channel of the bat 40S ribosome and its interacting partners uS5 (blue), uS3 (light green), eS30 (yellow), and helix 18 (h18) of the 18S rRNA (purple). (C) Key interactions of nsp1 CTD with rRNA, uS5, and uS3. Polar interactions are shown as black dotted lines. Select residues are rendered with semi-transparent density. (D) Interaction map of the SARS-CoV-2 nsp1 CTD with the 40S ribosome. The CTD residues are connected to key ribosomal interactors through dotted lines.

### Molecular basis of SARS-CoV-2 nsp1-mediated bat translation inhibition

Our cryo-EM structure resolves the nsp1 CTD embedded in the mRNA entry channel of the ribosome, confirming successful complex formation and enabling visualization of nsp1 recognition of the Rhileki 40S ribosome (**Fig 2B**). As seen in cryo-EM structures of nsp1 bound to the human ribosome, the CTD forms two alpha helices (consisting of residues 153-160 and 166-178) and interacts extensively with uS5, uS3, eS30, and helix h18 of the 18S rRNA (**Fig 2C-D**). Extensive van der Waals interactions are formed between nsp1 and uS5 involving nsp1 residues F157, W161, L173, and L177, and uS5 residues V106, P111, F124, V157, I151, and I190, along with aromatic stacking of nsp1 F157 and uS5 F124. Nsp1 helix 1 residues interact with uS3 through predominantly polar contacts with D152, E155, and E159 forming salt bridges with uS3 residues R116, R143, and K148, respectively. The 18S rRNA h18 forms multiple hydrogen bonds and electrostatic interactions with nsp1 helix 2 comprising residues 164-179. The phosphate groups of G606, G625, and U630 respectively interact with the positively charged side chains of R175, K164, and H165 whereas S167 is hydrogen-bonded to U630 and the G600 ribose. The conserved K164/H165 motif forms multiple contacts with U607, C624, G625, and U630, highlighting its importance in promoting CTD interactions within the bat ribosome, thereby rationalizing the loss of function of the nsp1 K164A/H165A double mutant (**Fig 1B-C**). Overall, nsp1 buries 1655 Å^2^ of its surface at the interface with the 40S, showcasing very extensive interactions despite its relatively small size, concurring with its high binding affinity^32,53^. Collectively, our data show that the nsp1 CTD interacts with the bat ribosome using a conserved binding mode to that observed for the human ribosome^21,24,25^, leading to translation inhibition via blockade of the mRNA entry channel.

### Nsp1 inhibits translation in a broad range of SARS-CoV-2 hosts

Given the observed nsp1 targeting of a conserved site within the Rhileki and human 40S ribosomes, we set out to understand the breadth of its inhibitory activity across a wide range of hosts known to be infected by SARS-CoV-2, including cats, dogs, minks and deers^33–35^. We therefore extended the transfection assay described in Figure 1A to other animal cell lines, including Crandell-Rees feline kidney epithelial cells (CRFK), canine fibroblasts cells (A72), American mink epithelial cells (Mv 1 Lu), Southern red muntjac (deer) fibroblast cells (Muntjac), and pig testis fibroblast cells (ST). All these mammals are susceptible to SARS-CoV-2 infection except pigs which we included in our panel to assess nsp1 activity in a species known to not be permissive to SARS-CoV-2^32–34,39^. Nsp1 mediated potent and concentration-dependent translation inhibition in CRFK, A72, Mv 1 Lu, Muntjac, and ST whereas the nsp1 K164A/H165A double mutant did not **(Fig 3, Fig S1)**. Differences in GFP and nsp1 expression levels across cell lines reflect the extent of nsp1-mediated translation inhibition, as seen most prominently in Muntjac cells which have relatively lower GFP and nsp1 expression that correlates to the lower nsp1 inhibition at 0.01 ng nsp1 (**Fig S1, Fig 3B**). These results demonstrate that nsp1 inhibits protein translation in a wide range of mammalian hosts, concurring with the strict conservation of its binding site on the ribosome across mammals.

**Figure 3.**
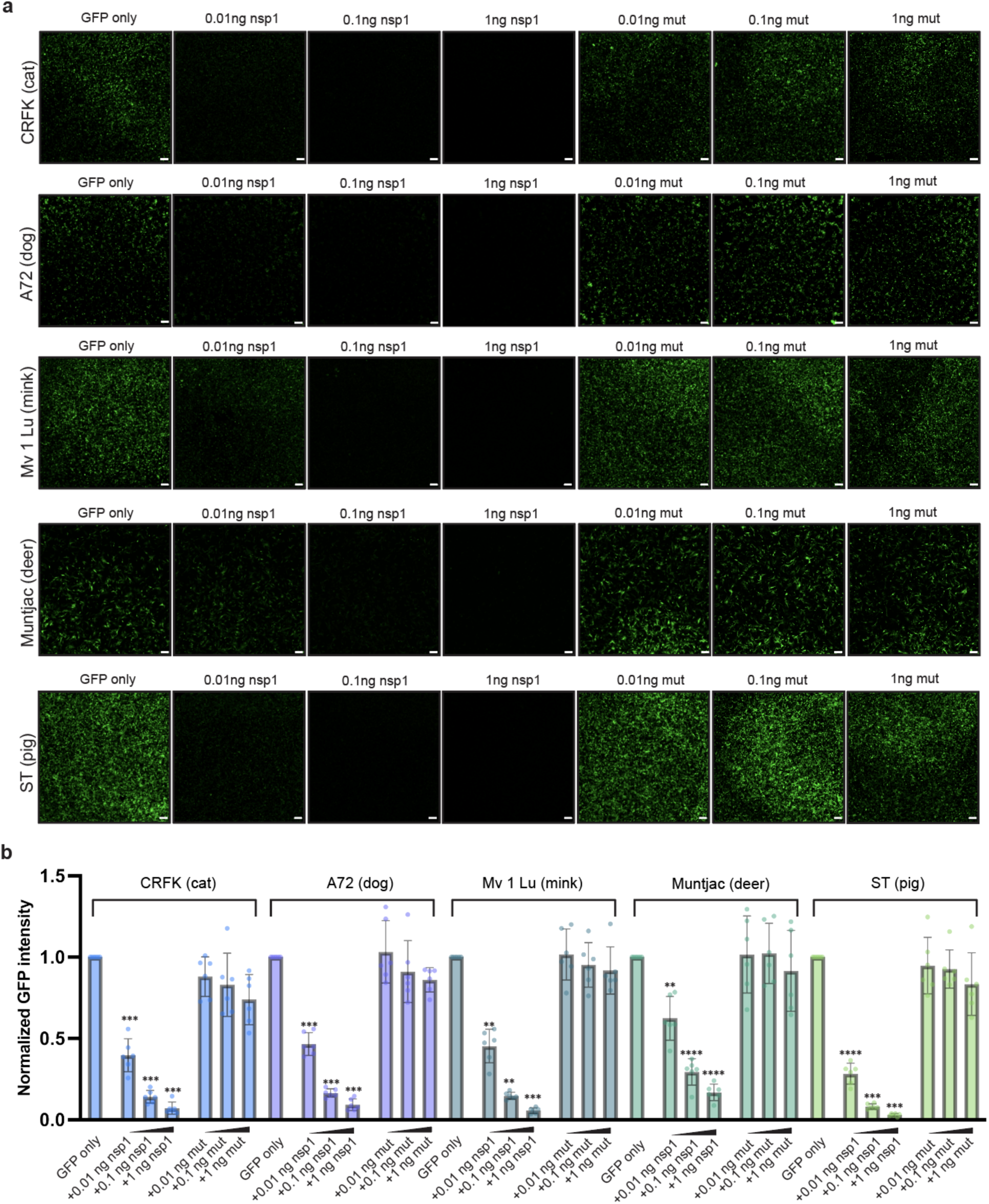
SARS-CoV-2 nsp1 inhibits translation in a wide range of mammalian species. (A) Live-cell fluorescence imaging of GFP expression in the presence (or absence) of varying amounts of wildtype nsp1 or of the K164A/H165A nsp1 mutant (mut) mRNA in Crandell-Rees feline kidney epithelial cells (CRFK), canine fibroblasts cells (A72), American mink epithelial cells (Mv 1 Lu), Southern red muntjac deer fibroblast cells (Muntjac), and pig testis fibroblast cells (ST). Scale bar: 100 µm. (B) Quantification of the nsp1-mediated dose-dependent inhibition of translation based on normalized GFP fluorescence intensity using the entire field of view. Bars represent the mean of six biological replicates shown as individual points. Each biological replicate is a mean of six technical replicates. Transfection with wildtype and mutant nsp1 mRNAs were compared to the GFP-only control using one-way ANOVA and follow-up Dunnet’s T3 multiple comparisons tests. See Fig S1 for GFP expression levels. P-values reported are adjusted for multiplicity. **: p<0.01, ***: p<0.005, ****: p<0.001.

## Discussion

Establishment of a successful viral infection depends on multiple factors, including viral entry, innate immune antagonism and the ability to hijack cellular machineries for replication. As a result, studies of non-structural proteins involved in innate immune antagonism are key to understanding host species tropism, spillover and pathogenicity. For instance, the Nipah virus and Hendra virus V and W proteins, which result from frameshifting of P transcripts, inhibit interferon signaling responses with the V proteins also being a determinant of pathogenicity in vivo^54–60^. The lack of V and W proteins in the related Cedar virus renders it unable to mediate similar innate immune antagonism which has been proposed to contribute to the low pathogenicity of this virus^61^.

Our results unveil the architecture of a bat 40S ribosome, which is extraordinarily conserved with its human counterpart despite extensive evolutionary divergence between the Primate and Chiropteran orders. In line with this result, SARS-CoV-2 nsp1 inhibits translation in bat cells and several other cells from cats, dogs, minks, deers and pigs by obstructing a highly conserved site in the mRNA entry channel, providing a molecular rationale for the broadly inhibitory nsp1 activity in several mammalian species. Nsp1 has also evolved to interfere with host mRNA in various ways such as prevention of host mRNA export from the nucleus^62^, induction of host mRNA cleavage^63^ and promoting evasion of direct Natural Killer cell cytotoxicity^64^. The multipronged role of nsp1 in immune evasion showcases the evolutionary arms race between viruses and hosts and the importance of nsp1 as a crucial factor in viral fitness for several coronaviruses^65–67^.

Nsp1 is one of the key players of the highly coordinated viral strategy to antagonize the host innate immune response by suppressing translation of antiviral genes such as interferons and NF-kB^68^. SARS-CoV-2-mediated innate immune antagonism also relies on several other non-structural proteins, such as ORF6 which blocks nuclear import and export^69^, rendering host cells incapable of responding to SARS-CoV-2 infection. As nsp1 efficiently inhibits translation in bat cells, the downstream effects of innate immune suppression are expected to be similar between bats and humans due to nsp1 targeting a conserved pathway. Future studies will investigate the activity of other SARS-CoV-2 and sarbecovirus innate immune modulators in bats to further our understanding of the mechanisms underlying immune tolerance of pathogenic sarbecoviruses.

Numerous viruses hinder host protein translation to selectively promote production of viral proteins through various strategies. At the transcriptional level, a multitude of viruses encode proteins that disrupt RNA polymerase II^70–72^ whereas others block the nuclear mRNA export machinery^62,73^. Targeting cellular mRNAs for degradation^44,74^ or preventing them from being translated through direct interference with translation factors, such as the eIF4F complex and poly(A)-binding protein, are other examples of strategies employed by viruses^75,76^. Despite the variety of approaches evolved to interfere with host protein translation, obstruction of the 40S ribosome mRNA channel is unique to nsp1^77^. Some alpha-CoV nsp1s also interact with the 40S ribosome to suppress host translation^78^ through a presumably distinct mechanism of action due to their lack of CTD. Conversely, host proteins have evolved to perform a similar role to nsp1. For instance, SERPINE mRNA-binding protein 1 (SERBP1) and tumor suppressor Pdcd4 are cellular factors binding to the mRNA entry channel and obstructing it to regulate translation initiation^79,80^. These various pathways illustrate the plethora of molecular solutions that evolved to control protein translation in physiological and pathological processes.

The continued circulation and antigenic evolution of SARS-CoV-2 is associated with successive waves of infection globally, imposing considerable stress on public health infrastructures^81–85^. The ability of nsp1 to mediate translation inhibition in a broad spectrum of mammalian species shows that it is not a limiting factor for SARS-CoV-2 spillovers to other species and brings up the potential for further spread of SARS-CoV-2 into new hosts, as previously observed in white-tailed deers and farmed minks^34,86^. The conserved nsp1-binding site on the 40S ribosome along with the reduced nsp1 genetic drift among SARS-CoV-2 variants^87^, relative to the spike glycoprotein, suggest the possibility of targeting this key virulence factor to develop next-generation COVID-19 countermeasures.

## Acknowledgements

We thank Eileen Chen and Gavin JD Smith from Duke-NUS for sharing the Rhileki cell line. This study was supported by the National Institute of Allergy and Infectious Diseases (DP1AI158186 and 75N93022C00036 to D.V.), an Investigators in the Pathogenesis of Infectious Disease Awards from the Burroughs Wellcome Fund (D.V.), a Shurl and Kay Curci Foundation Graduate Scholarship Award (to R.G.), the University of Washington Arnold and Mabel Beckman cryoEM center and the National Institute of Health grant S10OD032290 (to D.V.). D.V. is an Investigator of the Howard Hughes Medical Institute and the Hans Neurath Endowed Chair in Biochemistry at the University of Washington.

## Contributions

RG and DV designed the project. RG purified nsp1, ribosome and RNA samples used in this study. MC assisted with ribosome purification under the supervision of ARS and MGC. CL purified eIF1. RG and DA carried out cryoEM sample vitrification, data collection and processing. DA and YJP assisted with cryoEM analysis. RG and DV built and refined the atomic models. AA carried out transcriptomic analysis. RG and AA sequenced the *R. lepidus* 18S rRNA. RG carried out translation inhibition assays. RG and DV analyzed the data and wrote the manuscript with input from all authors.

## Methods

### Cell Culture and Reagents

Cell lines used in this study were HEK293T (Female cell line, CRL-3216), Rhileki (Unsexed)^88^, CRFK (Female cell line, CCL-94), Indian Muntjac (Male cell line, CCL-157), ST (Male cell line, CRL-1746), A-72 (Female cell line, CRL-1542), and Mv 1 Lu (Male and female mixed cell line, CCL-64). HEK293T, A-72, and CRFK were cultured in 10% FBS (Fisher Scientific-Cytiva), 1% penicillin-streptomycin (Thermo Fisher Scientific) in Dulbecco’s Modified Eagle Medium (DMEM) at 37°C, 5% CO_2_. Rhileki was cultured in 10% FBS, 1% penicillin-streptomycin, 1% sodium pyruvate (Thermo Fisher Scientific), and 1% non-essential amino acids (Thermo Fisher Scientific) in DMEM. ST and Mv 1 Lu were cultured in 10% FBS in Eagle’s Minimum Essential Medium (EMEM). Indian Muntjac was cultured in 20% FBS in Hamster’s F-10 Nutrient Mix (Thermo Fisher Scientific).

### Construct Design

pMCSG53 was used for cloning in *E. coli* DH10 cells, and for recombinant protein expression in *E. coli* BL21 (DE3) cells. SARS-CoV-2 Nsp1 full length (NC_045512.2) was cloned into pMCSG53 with a 6xHis and TEV (Tobacco Etch Virus) tag inserted before the N terminal domain for purification and cleavage.

eGFP transcript was obtained from Genscript including a 5’ cap, synthetic nonviral 5’ UTR (sequence: AAATAAGAGAGAAAAGAAGAGTAAGAAGAAATATAAGA), and 100 nt polyA tail. Nsp1 WT and mutant transcripts were obtained from Trilink with a 5’ cap, AGG initiation codon followed by genomic SARS-CoV-2 5’ UTR, 3XFLAG tag, and 80 nt polyA tail. All transcripts include N1-methyl pseudouridines in substitute of uridines.

### Bacterial Expression and Purification of Recombinant nsp1

Nsp1 was transformed and expressed in Escherichia coli BL21 (DE3) cells. Cells were grown at 37°C until the OD_600_ reached 0.6, when the cells were then induced by 1.0 mM isopropyl β-D-1-thiogalactopyranoside (IPTG) and shifted to incubation at 18°C for 18 hours in Luria’s Broth in the presence of ampicillin. Cells were harvested via centrifugation at 7000xg for 30 min, resuspended in lysis buffer [20 mM HEPES-NaOH pH 8.0, 500 mM NaCl, 5 mM MgCl2, 10 mM imidazole, 10% glycerol, 0.5 mM TCEP], and lysed using a C5 Emulsiflex. The lysate was clarified via centrifugation at 11,000xg for 40 min and filtered through a 0.45 µm membrane before being applied to a HisTrap HP column (Cytiva). Proteins were washed in lysis buffer supplemented with 20 mM imidazole before being eluted in lysis buffer supplemented with 300 mM imidazole. TEV cleavage was performed immediately after in TEV reaction buffer [50 mM Tris-HCl pH 8.0, 0.5 mM EDTA, 1mM DTT, 5% glycerol] with TEV protease (1 uL per 30 ug protein) and incubated at 30°C for 1 hour. SDS-PAGE gel was run to analyze cleavage products. Concentration of Nsp1 was performed in Amicon Ultra-15 Centrifugal Filter Concentrators with a 10 kDa molecular weight cutoff. Size exclusion chromatography (HiLoad Superdex 75, GE healthcare) was performed at 4°C in storage buffer [20 mM HEPES-NaOH pH 7.5, 500 mM NaCl, 5 mM MgCl2, 10% glycerol, 1 mM TCEP]. Purity of the proteins were analyzed by SDS-PAGE after each step.

### Bacterial Expression and Purification of Recombinant eIF1

Human eIF1 was purified as described previously^53,89^. Briefly, recombinant human eIF1 was expressed in *E. coli*, cleaved using recombinant TEV protease, and purified via ion-exchange chromatography.

### Purification of 40S ribosomal subunits

40S ribosomal subunits were purified from Rhileki cells following previously established protocols (Khatter et al., 2014) Rhileki cells were trypsinized to detach from cell culture flasks and pelleted via centrifugation at 1200 rpm for 5 min. The cell pellet was resuspended in lysis buffer [30 mM Tris-HCl pH 7.5, 300 mM NaCl, 30 mM MgCl2, 2% (v/v) Triton-X 100, 4 mM DTT] supplemented with RNAsin (Promega) and Halt protease inhibitor (Thermo) and incubated on ice for 10 min. Following brief centrifugation at 10,000 rpm for 10 min at 4°C, the lysate was loaded on a 30% sucrose cushion [20 mM Tris-HCl pH 7.5, 500 mM KCl, 30% (v/v) sucrose, 10 mM MgCl2, 0.1 mM EDTA pH 8.0, and 2 mM DTT] and pelleted via ultracentrifugation for 18 hours at 30,000 rpm in a Beckman Ti60 rotor at 4°C. Ribosomes were then resuspended in resuspension buffer [20 mM Tris-HCl pH 7.5, 500 mM KCl, 7.5% v/v sucrose, 2 mM MgCl2, 75 mM NH4Cl, 2mM puromycin, 2 mM DTT] supplemented with RNAsin and Halt protease inhibitor and incubated at 4°C for 1 hour, followed by 37°C for 1.5 hours. The resuspended ribosomes were then loaded on a 11% to 25% sucrose gradient [20 mM Tris-HCl pH 7.5, 500 mM KCl, 11%-25% (v/v) sucrose, 6 mM MgCl2] using a Beckman SW 32 Ti rotor at 17,000 rpm for 16 hours at 4°C. Sucrose gradient was prepared using the Biocomp gradient master. Fractions were collected by the Biocomp piston fractionator and fractions containing 40S subunits were combined, and concentrated using the Amicon Ultra-15 Centrifugal Filter Concentrators with a 100 kDa cutoff while being buffer exchanged into storage buffer [30 mM HEPES-KOH pH 7.4, 100 mM KOAc, 5 mM Mg(OAc2), 6% (v/v) sucrose, 2mM DTT]. After negative stain and 1% bleach agarose gels were used to confirm sample purity, small aliquots were flash frozen in liquid nitrogen and stored at -80°C.

### Transfection Assay and Western Blot Analysis of GFP Expression

Day before transfections, cells were seeded in a black 96-well glass bottom plate (Cellvis). The next day, 200 ng eGFP mRNA alone or eGFP with 0.01 ng, 0.1 ng, or 1 ng nsp1/nsp1 mutant were mixed with Lipofectamine 2000 in a 1ug:3uL ratio of mRNA to Lipofectamine and added to each well for six technical replicates. The cells were incubated with the transfection mixture for 16 hours in a 37°C, 5% CO_2_ incubator. Right before measurements, media was removed from the cells and replaced with pre-warmed FluoroBrite DMEM. Fluorescence was detected and quantified using the Biomek Cytation 5 with a GFP filter. The 16-bit raw images were processed in ImageJ to adjust the color display to green and a maximum value of 37870 was set for all images.

For GFP expression western blot analysis, after GFP quantification, cells were washed with ice-cold 1X phosphate buffered saline (PBS, pH 7.4) before adding 10 uL RIPA Lysis and Extraction buffer (Thermo) supplemented with SIGMAFAST™ Protease Inhibitor Cocktail Tablets (Sigma-Aldrich) to each well and incubated for 5 min. The cell lysate was collected and centrifuged for 15 min at 14,000xg at 4°C. The supernatant was collected and combined with 4X SDS-PAGE Sample Loading Buffer and boiled at 95°C for 5 min. 8 uL of each transfection sample (along with non-transfected negative control) was run through a 4%–20% gradient Tris-Glycine gel (BioRad) and transferred to a PVDF membrane using the mixed molecular weight protocol of the Trans-Blot Turbo System (BioRad). The membrane was blocked with 5% milk in TBS-T (20 mM Tris-HCl pH 8.0, 150 mM NaCl) supplemented with 0.05% Tween-20 at room temperature and with agitation. After 1 h, the GFP monoclonal antibody (1:3000 dilution, GF28R, Thermo Fisher) and incubated for 1h at room temperature with agitation. The membrane was washed three times with TBS-T and an IRDye 680 RD Goat anti-Mouse IgG secondary antibody (1:20,000 dilution, LI-COR, 926-68070) was added and incubated for 1 h at room temperature. The membrane was washed three times with TBS-T after which a LI-COR processor was used to develop the western blot.

Membranes in figures were cropped to show area of interest, with full uncropped image scans provided in the extended data file.

### Transfection Assay and Western Blot Analysis of Nsp1 Expression

Day before transfections, cells were seeded in a 12-well cell culture plate (Genesee Scientific) and incubated overnight. The next day, 400 ng, 200 ng, and 100 ng of nsp1 and nsp1 mutant were transfected into each well with Lipofectamine 2000 in a 1ug:3uL ratio of mRNA to Lipofectamine and incubated overnight. After 16 hours, cells were washed twice with ice-cold 1X phosphate buffered saline (PBS, pH 7.4) before adding 50 uL RIPA Lysis and Extraction buffer (Thermo) supplemented with SIGMAFAST™ Protease Inhibitor Cocktail Tablets (Sigma-Aldrich) for 5 min. The cell lysate was collected and centrifuged for 15 min at 14,000xg at 4°C. The supernatant was collected and combined with 4X SDS-PAGE Sample Loading Buffer and boiled at 95°C for 5 min.

For nsp1 expression western blot analysis, 8 uL of each transfection sample (along with non-transfected negative control) was run through a 4%–20% gradient Tris-Glycine gel (BioRad) and transferred to a PVDF membrane using the mixed molecular weight protocol of the Trans-Blot Turbo System (BioRad). The membrane was blocked with 5% milk in TBS-T (20 mM Tris-HCl pH 8.0, 150 mM NaCl) supplemented with 0.05% Tween-20 at room temperature and with agitation. After 1 h, the nsp1 polyclonal antibody (1:1000 dilution, SARS-CoV-2 Nsp1 Antibody, Thermo Fisher PA5-11694) and incubated overnight at 4°C with agitation. The next day, the membrane was washed three times with TBS-T and an IRDye 680RD Goat anti-Rabbit IgG secondary antibody (1:20,000 dilution, LI-COR, 926-68073) was added and incubated for 1 h at room temperature. The membrane was washed three times with TBS-T after which a LI-COR processor was used to develop the western blot.

### PCR amplification of 18S ribosomal RNA and RPS10

One day before extraction, Rhileki cells were plated on two 100 mm cell culture plates (Corning) and incubated overnight in 37°C, 5% CO_2_. The next day, cells were washed with ice-cold 1X PBS and total RNA was extracted from Rhileki cells using the Quick-RNA Miniprep Kit (Zymo Research) following the recommended protocol. Extracted RNA was quantified and stored at -80°C. For PCR amplification of ribosomal protein S10 and 18S rRNA, primers were designed based on the *Rhinolophus ferrumequinum* sequences (GCF_004115265.2) with forward primer ACCTGGTTGATCCTGCCAGTAG and reverse primer TAATGATCCTTCCGCAGGTTCACC for 18S, and forward primer CCGGGACGYMGAAGTCGTAA and reverse primer TGCAAAGAYGGGCACACAAG for RPS10. cDNA was subsequently synthesized using the Lunascript RT Supermix (NEB) with the recommended protocol. PCR reactions were performed with KAPA HiFi HotStart ReadyMix (Roche) and CloneAmp HiFi PCR Premix (Takara Bio). PCR products were run on 1% Tris acetate-EDTA (TAE) agarose gels and extracted using the Zymoclean Gel DNA Recovery Kit (Zymo). Products were sent for sequencing for confirmation of the correct amplicon.

### Transcriptome sequencing sample preparation and data analysis

Total RNA from Rhileki cells was extracted as described above. Sequencing libraries were prepared using the TruSeq Stranded mRNA kit (Illumina) and sequenced on a 2 x 150 bp run on an Illumina NovaseqX+ by Psomagen. A total of 116,395,242 sequencing reads were obtained for the sample. The sequencing reads were quality and adapter trimmed using Trimmomatic v0.39 and aligned to the homologous ribosomal transcripts of *R. ferrumequinum* (RefSeq accession number: GCF_004115265.2) using Geneious Prime. The sequence alignments were visualized in Geneious Prime and the final sequences of the *R. lepidus* ribosomal transcripts were manually called. Sequencing reads are available under the NCBI Bioproject PRJNA1190498 and ribosomal coding sequences are included in supplementary materials.

### Cryo-EM Sample Preparation and Data Collection

Nsp1 bound Rhileki 40S ribosome complex was prepared by mixing a 1:5:5 molar ratio of 40S ribosome:nsp1:eIF1 followed by a 15 minute incubation at room temperature and kept on ice for the remaining time until grids were frozen. 3 uL of 80 nM complex was applied onto fresh glow discharged R 2/2 UltrAuFoil grids prior to plunge freezing using a vitrobot MarkIV (ThermoFisher Scientific) with a blot force of -1, 6 sec blot time, and 40 sec waiting time at 100% humidity and 22°C. The data was acquired using an FEI Titan Krios transmission electron microscope operated at 300 kV and equipped with a Gatan K3 direct detector and Gatan Quantum GIF energy filter, operated in zero-loss mode with a slit of 20 eV. Automated data collection was carried out using SerialEM^90^ at a nominal magnification of 105,000x with a pixel size of 0.829 Å. A total of 14,216 micrographs were collected with a defocus range between -0.6 and -1.6 μm with stage tilt angle of 0° and 30°. Movie frame alignment, estimation of the microscope contrast-transfer function parameters, particle picking, and extraction were carried out using cryoSPARC Live^91^, and all downstream processing steps were done in cryoSPARC^92^. Particles were extracted with a box size of 432 pixels and downsampled by a factor of 6. One round of reference-free 2D classification was performed to select well-defined particle images. Initial model generation was performed using ab-initio reconstruction and the resulting map was used for non-uniform refinement. The resulting map was used to re-extract 1,205,667 particles with extraction box size of 512 pixels cropped to 384 pixels. The re-extracted particles were used for subsequent non-uniform refinement with per-particle defocus and global CTF refinement. This was followed by reference-based motion correction and subsequent non-uniform refinement with per-particle defocus, global CTF refinement, and Ewald sphere negative correction to obtain the consensus map at 2.1 Å (EMDB XXXX). Masks for the 40S head and body were created in UCSF ChimeraX^93^ and low pass filtered to 20Å in cryoSPARC. Local refinement for the head and body regions were conducted separately using the same consensus map in order to yield final reconstructions of the 40S head and body maps at 2.1Å (EMDB XXXX and EMDB XXXX. Reported resolutions are based on the gold-standard Fourier shell correlation (FSC) of 0.143 criterion. See also Table S1 and Fig S3.

### Cryo-EM Data Processing and Model Building

Initial model building used ModelAngelo^94^ with the Rhileki 40S - nsp1 consensus map and sequences for nsp1, eIF1, and the 40S ribosome (**Data S1**). The structure was then manually rebuilt and extended in the locally refined head and body maps using Coot^95^ and refined using Phenix^96^, ISOLDE^97^ and Rosetta^98^. Human 40S structures were used to assist the process (PDB: 8PPK)^31^ **(Table S2)**. Validation used Molprobity^99^ and Phenix. Modified residues and nucleotides were included according to density and using human 80S ribosome (PDB: 8QOI)^100^ and human 40S ribosome (PDB: 8PPK) as guides. Pseudouridines were added to the model when justified by the presence of a water molecule next to the N1 position of the base^100^ . Structures are deposited in the PDB with accession codes XXXX, XXXX, and XXXX for the consensus (excluding head region except RPS3), body, and head structures respectively.

### Statistical Analysis

Transfection assays were conducted 6 times with 6 biological repeats. Data were presented by means ± SD as indicated in the figure legends. One-way ANOVA and Dunnett’s T3 multiple comparison tests were conducted for all statistical analyses using GraphPad Prism 8 unless specified otherwise. *P* < 0.05 was considered significant. \**P* < 0.05,\*\**P* < 0.01, \*\*\**P* < 0.005, and \*\*\*\**P* < 0.001.

## Supplementary Materials

**Supplementary Figure S1.**
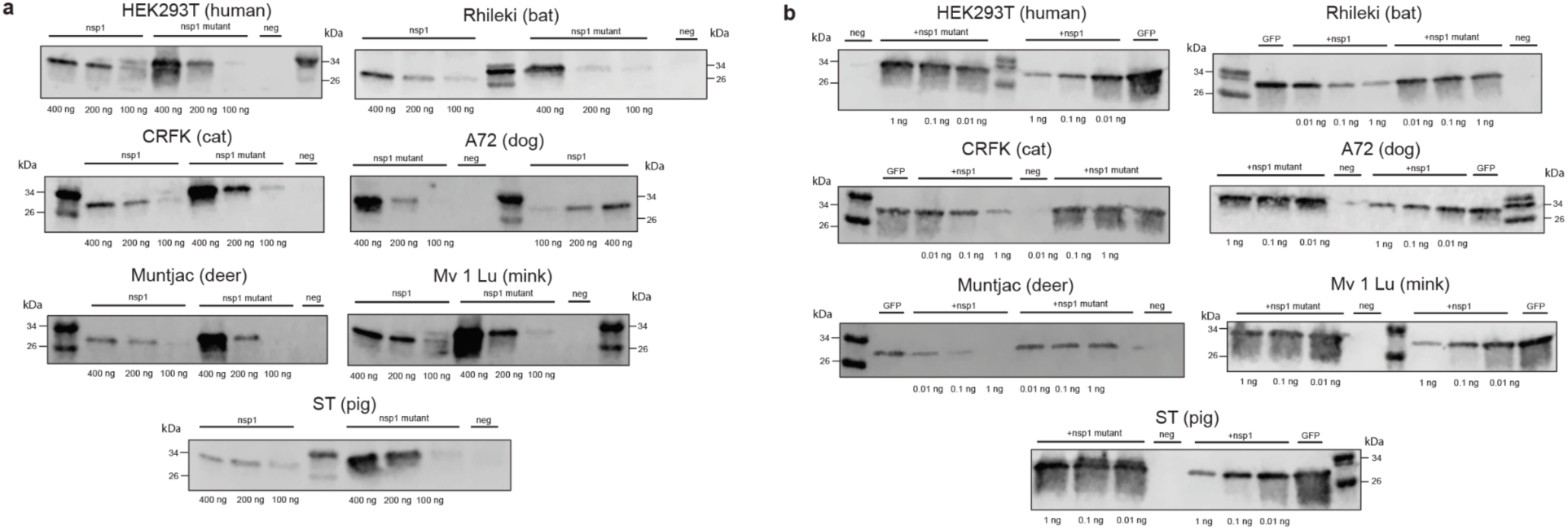
Protein expression in animal cell lines, related to Figures 1 and 3. (a) Western blots of nsp1 and nsp1 mutant expression using anti-nsp1 polyclonal antibody (Thermo) in seven different animal cell lines with transfection amounts listed below the images. Transfection assays for these western blots were performed in 12-well plates with increased amounts of mRNA to enable detection of expression of the wildtype and of the K164A/H165A mutant nsp1 constructs, which could not be detected in a 96-well format (with much lower amounts of mRNA transfected) used for Figures 1 and 3. (b) Western blot analysis of GFP expression in the presence or absence of the wildtype or the K164A/H165A mutant nsp1 shown in Figures 1 and 3 using anti-GFP monoclonal antibody GF28R (Thermo). Neg: negative control.

**Supplementary Figure S2.**
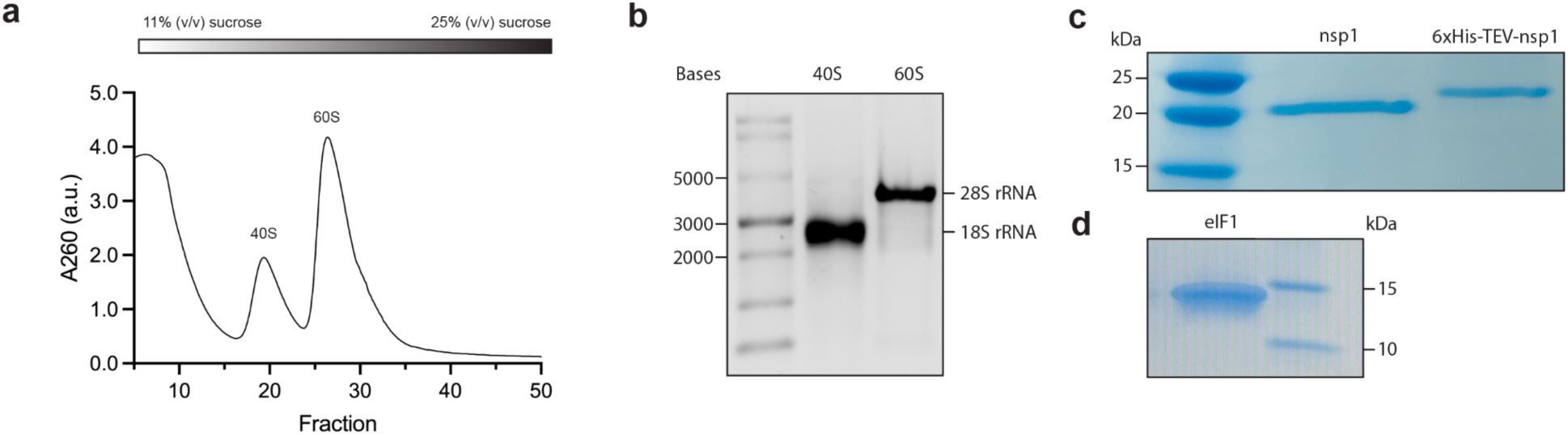
Purification of the bat Rhileki ribosome and SARS-CoV-2 nsp1, related to Figure 2. (a) Isolation of the Rhileki ribosomal 40S and 60S subunits using a 11-25% (v/v) sucrose density gradient. The purification process was followed using absorbance at 260 nm (A_260_) of each fraction. (b) 1% bleach agarose gel showing separation of the 40S and 60S ribosomal subunits through 18S and 28S rRNA respectively. An RNA ladder was used to deduce separation of the 18S and 28S bands. (c) SDS-PAGE gel of recombinant SARS-CoV-2 nsp1 with or without the 6xHis-TEV tag. (d) SDS-PAGE gel of recombinant human eukaryotic initiation factor 1 (eIF1). TEV: Tobacco Etch Virus protease cleavage site.

**Supplementary Figure S3.**
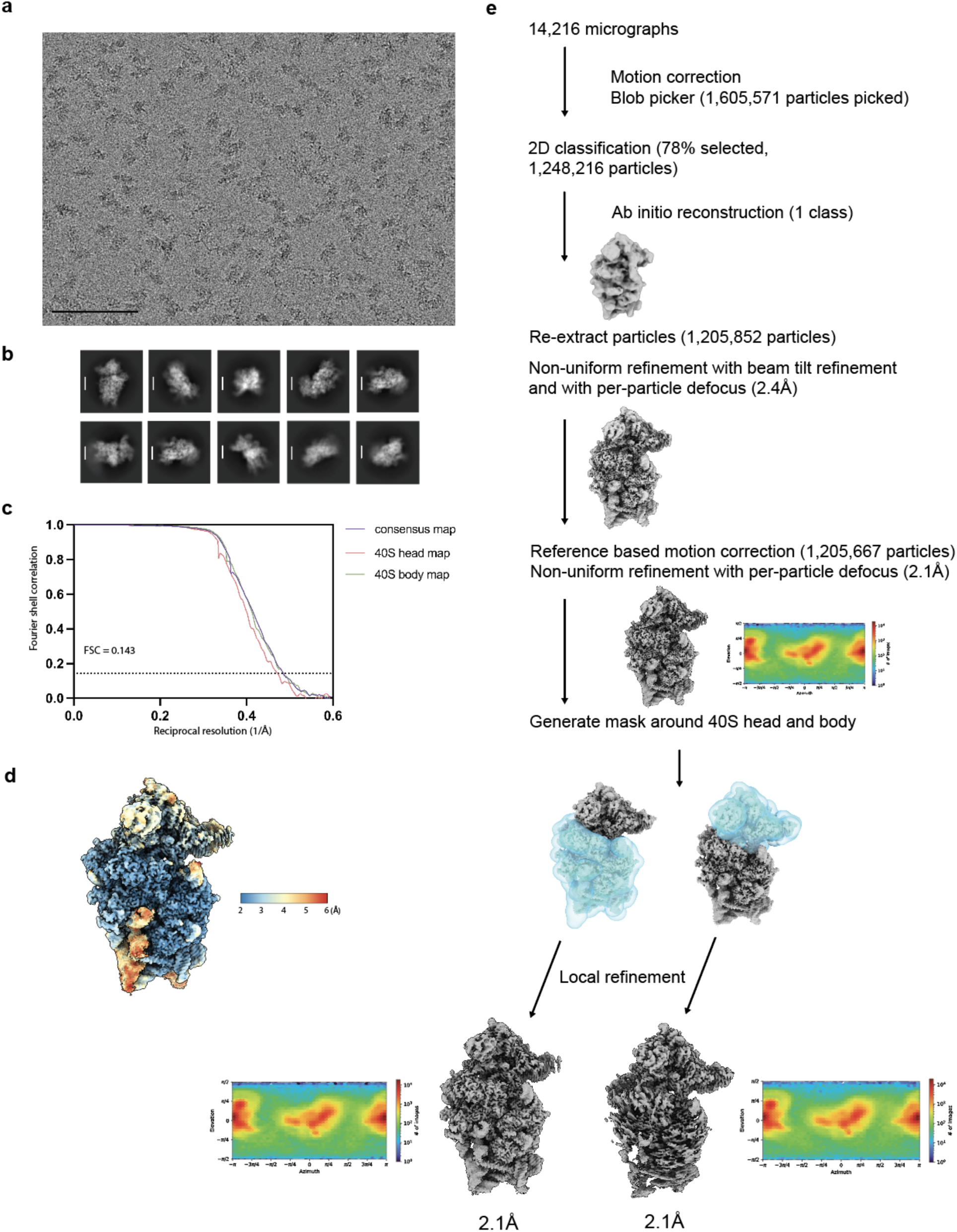
Cryo-EM workflow of SARS-CoV-2 nsp1 bound 40S Rhileki ribosome structure, related to Figure 2. (a) Representative electron micrograph and (b) 2D class averages. Scale bars of the micrograph and class averages are 100 nm and 100Å, respectively. (c) Combined gold-standard fourier shell correlation (FSC) curves for the consensus map (purple line) and the locally refined maps, separated for the 40S head (pink line) and body (green line). (d) Local resolution map of the consensus refinement. (e) Data processing flowchart for the reconstruction of the consensus map and the subsequent locally refined maps. Angular distribution plots with all the particles contributing to the final maps are shown for the three maps.

**Supplementary Figure S4.**
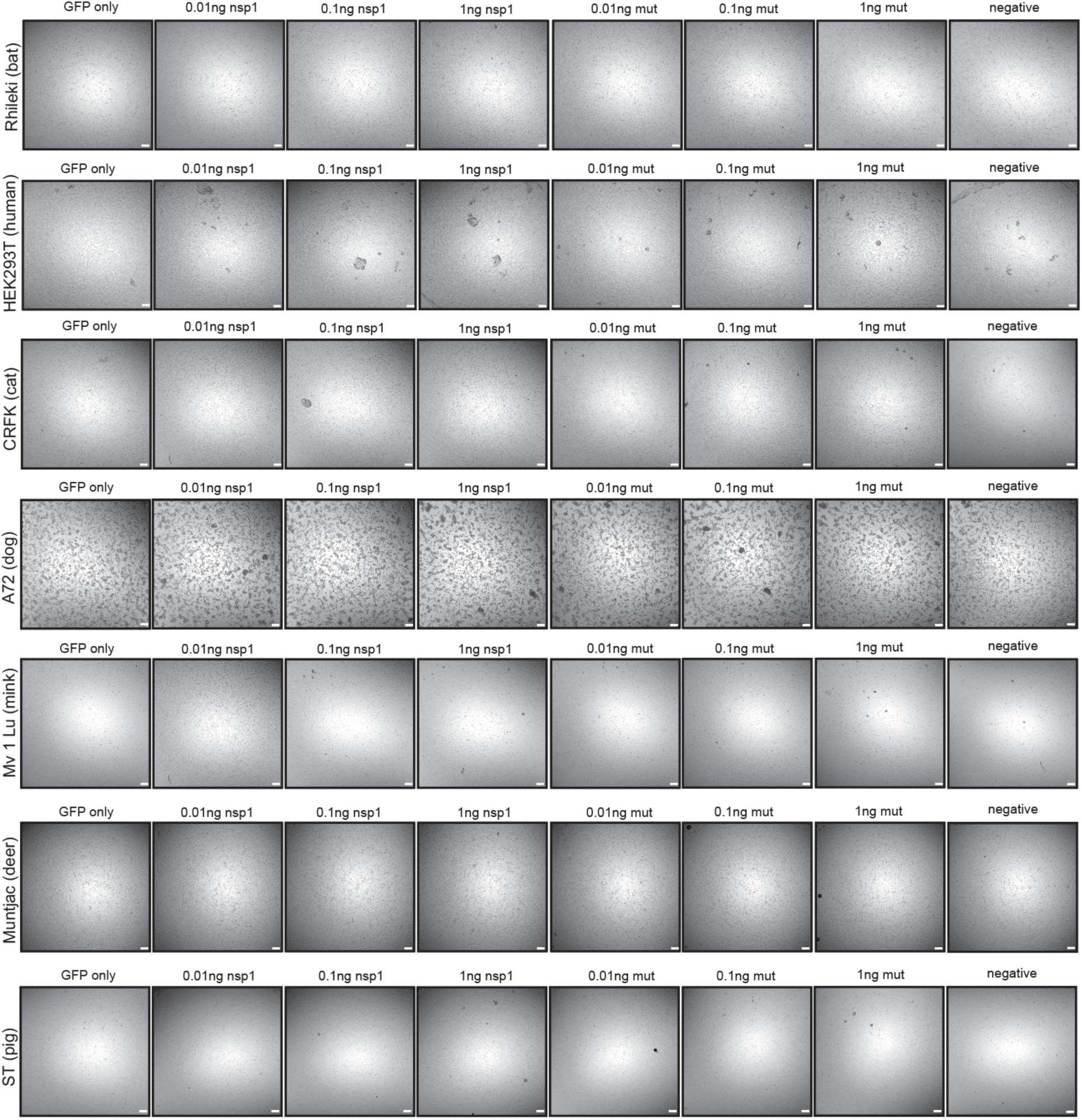
Bright-field images of cell lines used for transfections, related to Figures 1 and 3. Corresponding bright-field images of fluorescence images. Negative: non-transfected cells. Scale bar: 100 µm.

**Table S1:** Cryo-EM data collection, refinement, and validation statistics, related to Figure 2 and Figure S3.

|  | Rhileki ribosome<br>consensus PDB<br>### EMD ### | Rhileki ribosome<br>40S body PDB ###<br>EMD ### | Rhileki ribosome<br>40S head PDB ###<br>EMD ### |
| --- | --- | --- | --- |
| **Data collection and processing** |  |  |  |
| Magnification | 105,000 |  |  |
| Voltage (kV) | 300 |  |  |
| Electron exposure (e-/Å<sup>2</sup>) | 30.7 |  |  |
| Defocus range (µm) | -0.6 - -1.6 |  |  |
| Pixel size (Å) | 0.829 |  |  |
| Symmetry imposed | C1 |  |  |
| Initial particle images (no.) | 1,605,571 |  |  |
| Final particle images (no.) | 1,205,667 |  |  |
| Map resolution (Å) | 2.1 |  |  |
| FSC threshold | 0.143 |  |  |
| **Refinement** |  |  |  |
| Model resolution (Å) | 2.2 | 2.2 | 2.4 |
| FSC threshold | 0.5 | 0.5 | 0.5 |
| Map sharpening *B* factor (Å<sup>2</sup>) | 55.1 | 56.7 | 59.2 |
| **Model composition** |  |  |  |
| Non-hydrogen atoms | 52405 | 56113 | 25366 |
| Protein residues / Nucleotides | 3237 / 1218 | 3087 / 1281 | 1897 / 483 |
| Ligands | 72 | 83 | 32 |
| Water | 1639 | 1640 | 826 |
| ***B* factors (Å<sup>2</sup>)** |  |  |  |
| Protein | 82.66 | 89.4 | 46.3 |
| Nucleotide | 92.92 | 108.6 | 32.7 |
| Ligand | 76.84 | 83.6 | 29.0 |
| Water | 67.21 | 78.9 | 26.0 |
| **R.m.s. deviations** |  |  |  |
| Bond lengths (Å) | 0.003 | 0.006 | 0.003 |
| Bond angles (°) | 0.508 | 0.579 | 0.594 |
| **Validation** |  |  |  |
| **MolProbity score** | 1.26 | 1.37 | 1.55 |
| **Clashscore** | 4.95 | 6.00 | 5.64 |
| **Poor rotamers (%)** | 0.57 | 1.15 | 1.60 |
| **Ramachandran plot** |  |  |  |
| **Favored (%)** | 98.89 | 99.11 | 97.57 |
| **Allowed (%)** | 1.11 | 0.89 | 2.43 |
| **Disallowed (%)** | 0.00 | 0.00 | 0.00 |

**Table S2:** Model contents and sequences used for structure building, related to Figure S3.

| Name | Alternative Name | Chain ID | Length (residues) | Modeled residues | UniProt ID (proteins) |
| --- | --- | --- | --- | --- | --- |
| 40S Ribosomal RNA / Protein |  |  |  |  |  |
| 18S rRNA |  | i | 1869 | 1-232,287-689,745-751,792-1869 | NA |
| S2 | uS5 | A | 293 | 58-278 | NA |
| S3 | uS3 | B | 243 | 4-227 | NA |
| S3a | eS1 | C | 264 | 2-8,23-232 | NA |
| S4 | eS4 | D | 263 | 2-261 | NA |
| S5 | uS7 | E | 204 | 15-129,133-204 | NA |
| S6 | eS6 | F | 249 | 1-231 | NA |
| S7 | eS7 | G | 194 | 5-108,111-194 | NA |
| S8 | eS8 | H | 208 | 2-206 | NA |
| S9 | uS4 | I | 194 | 2-181 | NA |
| S10 | eS10 | J | 165 | 1-96 | NA |
| S11 | uS17 | K | 158 | 3-24,31-153 | NA |
| S12 | eS12 | K | 132 | 12-92, 102-128 | NA |
| S13 | uS15 | M | 151 | 2-151 | NA |
| S14 | uS11 | N | 151 | 18-151 | NA |
| S15 | uS19 | O | 145 | 13-136 | NA |
| S15a | uS8 | P | 130 | 2-130 | NA |
| S16 | uS9 | Q | 146 | 7-146 | NA |
| S17 | eS17 | R | 135 | 2-132 | NA |
| S18 | uS13 | S | 152 | 2-145 | NA |
| S19 | eS19 | T | 145 | 3-143 | NA |
| S20 | uS10 | U | 119 | 17-116 | NA |
| S21 | eS21 | V | 83 | 1-83 | NA |
| S23 | uS12 | W | 143 | 2-141 | NA |
| S24 | eS24 | X | 132 | 4-128 | NA |
| S25 | eS25 | Y | 125 | 42-112 | NA |
| S26 | eS26 | Z | 115 | 2-99 | NA |
| S27 | eS27 | a | 84 | 2-84 | NA |
| S27a | eS31 | b | 156 | 80-150 | NA |
| S28 | eS28 | c | 69 | 7-68 | NA |
| S29 | uS14 | d | 56 | 2-56 | NA |
| S30 | eS30 | e | 133 | 78-119, 126-132 | NA |
| L41 | eL41 | f | 25 | 1-22 | NA |
| RACK1 |  | g | 317 | 3-274,280-314 | NA |
| RPSA | LRP | h | 295 | 2-213 | NA |
| SARS-CoV-2 nsp1 |  | j | 180 | 148-180 | P0DTC1 |
| eIF1 |  | k | 113 | 30-48,54-76,81-100,110-112 | P41567 |

